# EpitopeVec: Linear Epitope Prediction Using Deep Protein Sequence Embeddings

**DOI:** 10.1101/2020.11.26.395830

**Authors:** Akash Bahai, Ehsaneddin Asgari, Mohammad R.K. Mofrad, Andreas Kloetgen, Alice C. McHardy

## Abstract

**Motivation:** B-cell epitopes (BCEs) play a pivotal role in the development of peptide vaccines, immunodiagnostic reagents, and antibody production, and thus generally in infectious disease prevention and diagnosis. Experimental methods used to determine BCEs are costly and time-consuming. It thus becomes essential to develop computational methods for the rapid identification of BCEs. Though several computational methods have been developed for this task, cross-testing of classifiers trained and tested on different datasets revealed their limitations, with accuracies of 51 to 53%.

**Results:** We describe a new method called EpitopeVec, which utilizes residue properties, modified antigenicity scales, and a Protvec representation of peptides for linear BCE prediction with machine learning techniques. Evaluating on several large and small data sets, as well as cross-testing demonstrated an improvement of the state-of-the-art performances in terms of accuracy and AUC. Predictive performance depended on the type of antigen (viral, bacterial, eukaryote, etc.). In view of that, we also trained our method on a large viral dataset to create a linear viral BCE predictor.

**Availablity:** The software is available at https://github.com/hzi-bifo/epitope-prediction under the GPL3.0 license.

**Supplementary information:** Supplementary data are available at *Bioinformatics* online.

## 1 Introduction

Antibodies are critical components of the humoral immune response that recognize and bind to antigens of pathogens such as bacteria and viruses (Janeway, 2012). The region of an antigen recognized by these antibodies is known as an epitope. Epitopes can either be a continuous stretch of amino acids within an antigen sequence (linear B-cell epitope) or amino acids potentially separated in the sequence, but located closely in 3D-protein structure (conformational B-cell epitope). Identification of B-cell epitopes (BCEs) is important for applications such as peptide-based vaccine design (Dudek *et al.*, 2010), immuno-diagnostic tests (Noya *et al.*, 2005), and synthetic antibody production (Hancock and O’Reilly, 2005). As experimental determination of BCEs is time consuming and expensive, computational prediction can play a pivotal role in the development of new vaccines and drugs against common viral pathogens such as human immunodeficiency virus, hepatitis, or influenza viruses (Pellequer *et al.*, 1991; Dudek *et al.*, 2010; Bryson *et al.*, 2010).

Although the majority of naturally occurring BCEs are conformational (Barlow *et al.*, 1986), prediction of linear BCEs has received much attention (Flower, 2007) as they are used for the synthesis of peptide-based vaccines. The earliest methods for epitope prediction evaluated only one physiochemical property of the constituent amino acids, such as surface accessibility (Emini *et al.*, 1985), flexibility (Karplus and Schulz, 1985), hydrophobicity (Levitt, 1976), or antigenicity (Kolaskar and Tongaonkar, 1990). Some of these methods that are still accessible online are: PREDITOP (Pellequer and Westhof, 1993), PEOPLE (Alix, 1999) and BEPITOPE (Odorico and Pellequer, 2003). These algorithms calculate the average amino acid propensity scale for individual features over a sliding window along the query protein sequence. If these predicted scales are above a certain cut-off for a continuous stretch on the protein then that region is determined to be a BCE. However, assessment of 484 propensity scales revealed that such scales are unreliable in detecting BCEs and barely outperformed random BCE selection when used based on a single amino acid feature or even a combination thereof (Blythe and Flower, 2009).

With the increased availability of experimentally identified epitopes, new methods were developed based on several propensity scales, including amino acid features that had not been included before (Yang and Yu, 2009). Such methods, employing Machine Learning (ML) approaches to distinguish BCEs from non-BCEs in the amino acid sequence, have shown better accuracy than single propensity scale based methods. For training, B-cell epitopes are presented as feature vectors derived from different amino acid properties such as propensity scales, amino acid composition, Amino Acid Pair (AAP) antigenicity scale (Chen *et al.*, 2007) or the Amino Acid Trimer (AAT) antigenicty scale (Yao *et al.*, 2012). Some examples of ML based methods for BCE prediction are BepiPred (Larsen *et al.*, 2006), ABCPred (Saha and Raghava, 2006), LBTope (Singh *et al.*, 2013), AAP (Chen *et al.*, 2007), BCPred (El-Manzalawy *et al.*, 2008) and SVMTrip (Yao et al., 2012). Most of these recent methods are trained on datasets compiled from databases of experimentally verified epitopes such as BciPep (Saha et al., 2005) or IEDB (Vita et al., 2009). A notable issue seems to be that all aforementioned methods lack high accuracy in a cross-testing approach, where ML training and testing are performed on independent datasets, which raises doubts about the generalization ability of all such approaches.

We here describe a method that combines commonly used propensity scales, residue features, modified antigenicity scales, and ProtVec (Asgari and Mofrad, 2015; Asgari et al., 2019b; Asgari, 2019) for vector representation of the peptides instead of the commonly used one-hot encoding method. ProtVec (Protein Vectors) is a distributed vector representation of proteins over k-mers (Asgari and Mofrad, 2015) trained using large protein databases (e.g., SwissProt or UniRef), using neural networks analogous to language-modeling (here in particular, a skip-gram neural network), which can predict the surrounding k-mers around a given k-mer. We use Support Vector Machines (SVM) and Random Forest (RF) as the predictive model and selected the SVM as our final model. We trained and tested our method on multiple small and large datasets derived from the BciPep and IEDB databases and compared the performance to the state-of-the-art methods. We also re-evaluated previously published methods on datasets on which they have not been tested before. We tested the reliability and generalization ability of our approach by cross-testing on datasets from different sources. Lastly, we trained our method on a large viral dataset to construct a predictor for linear viral BCEs.

## 2 Materials & Methods

### 2.1 Datasets

#### 2.1.1 BCPred dataset

We used BCPred (El-Manzalawy *et al.*, 2008) as our main training dataset, which was constructed from the Bcipep (Saha *et al.*, 2005) database. EL-Manzalawy et al. retrieved a set of 947 unique 20-mer epitopes and removed the redundant peptides by setting a homology cut-off of 80% to obtain a final set of 701 positive epitopes. A non-epitope (negative) set of 701 peptides was generated by randomly extracting 20-mer peptides from the SwissProt database while making sure that none of the negative peptides occurred in the epitope set.

#### 2.1.2 Chen dataset

This dataset was introduced by Chen et al. (Chen *et al.*, 2007) with the AAP scale. The epitope dataset was derived from the BCIPep database as well, while the non-epitope set was generated by randomly selecting 872 20-mer peptides from the Swiss-Prot database. However, the positive set contains redundant peptides as no homology cut-off was applied to this set unlike the BCPred dataset.

#### 2.1.3 ABCPred dataset

ABCPred (Saha and Raghava, 2006) was one of the first ML based methods for predicting linear BCEs. Saha et al. constructed datasets with epitope length ranging from 12 to 20-mers. We used the ABCPred16 dataset, which consists of 700 epitopes and 700 non-epitopes, as the original ABCPred method reported the highest performance on this particular dataset and we wanted to compare our method directly with ABCPred.

#### 2.1.4 Blind387 dataset

This dataset was also created by Saha et al. (Saha and Raghava, 2006) and consists of 187 epitopes and 200 non-epitopes. We used this dataset as an independent test set. The length of the peptides is not fixed here and it varies from 5 to 30 amino acids.

#### 2.1.5 LBTope dataset

LBTope (Singh et al., 2013) was the first method to use new datasets compiled from IEDB. There were datasets published with variable- and fixed-length epitopes. In this study we only used the LBTope_fixed_nr dataset which has a fixed length of 20 amino acid and is 80% non-homologous. The negative set was compiled from the protein sequences of the antigens while making sure that none of the peptides overlap with the positive set.

#### 2.1.6 Viral dataset

We downloaded viral peptides from IEDB that were reported as epitopes (positive) and similarly viral peptides tht were reported as non-epitopes (negative). The peptide length was not fixed and it varies from 6 to 46 amino acids. We used CD-HIT (Huang *et al.*, 2010) to remove homologous peptide sequences (cut-off of 80% for positive and 90% for negative). As the same peptide can be reported to be an epitope or a non-epitope for different neutralization experiments, we removed the peptides that are similar in both the sets using CD-HIT to finally obtain a dataset of 4432 peptides that are epitopes (positive) and 8460 peptides that are non-epitopes (negative).

### 2.2 Feature Representation of Peptides

#### 2.2.1 Amino Acid Composition

Amino acid composition (AAC) is a vector specifying the relative abundance of each amino acid in the peptide. It can be formulated as:

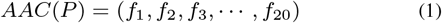

where 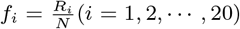 is the percentage of composition o: amino acid type *i, R_i_* is the count of type *i* in the peptide, and *N* is the peptide length.

#### 2.2.2 Dipeptide Composition

Dipeptide composition (DPC) is a vector specifying the abundance of dipeptides normalized by all possible dipeptide combinations. It has fixed length of 400 (20×20) features. It can be formulated as:

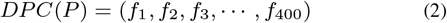

where 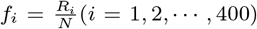 is the percentage of composition of dipeptide type *i, R_i_* is the count of type *i* in the peptide, and *N* is the peptide length.

#### 2.2.3 Chain-transition-distribution

Chain-transition-distribution (CTD) was introduced by Dubchak et al. (Dubchak et al., 1995) for predicting protein-folding classes and has been successfully applied for many protein sequence based classification problems (Zou et al., 2013). The standard amino acids are classified into three different groups: polar, neutral, and hydrophobic. Composition (C) is the number of amino acids of a particular property divided by the total number of amino acids. Transition (T) characterizes the percent frequency with which amino acids of a particular property is followed by amino acids of a different property. Distribution (D) measures the chain length within which the first, 25, 50, 75 and 100% of the amino acids of a particular property is located respectively. We used the PyDPI (Cao et al., 2013) package for calculating this feature.

#### 2.2.4 Amino Acid Pair antigenicity scale

The AAP antigenicity scale was introduced by Chen et al. in 2008. It is the ratio of occurrence frequency of amino acid pairs in the positive set compared to the negative set. We count the number of each type of dipeptide in the positive set and then calculate the frequency of each by dividing it by the total number of dipeptides in the entire positive set. Similarly, we do this for the negative dataset. The antigenicity value for each dipeptide is then the logarithm of the frequency in the positive set divided by the frequency in the negative set.

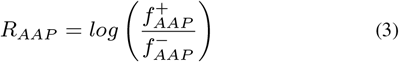

We normalize the scale between 1 and −1 to avoid the dominance of an individual propensity value. The normalization is defined as follows:

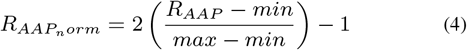

where max and min represent the maximum and minimum values among all the possible *R_AAP_* values before the normalization. For the positive set we use the BCIPEP dataset and for the negative dataset we chose the entire UniProt50 database from SwissProt (Bairoch, 2000) which contains 248,858 protein sequences.

#### 2.2.5 Amino Acid Trimer antigenicity scale

The AAT antigenicity scale was first introduced in SVMTrip (Yao et al., 2012) and it is similar to AAP scale except that in this we use amino acid triplets instead of amino acid pairs. Similarly, for the positive set we use the epitope dataset and for the negative dataset we choose UniProt50. The AAT scale is the logarithm of the ratio of the frequency of amino acid triplets in positive set to their frequency in the negative set.

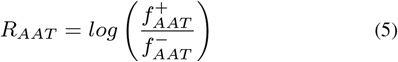

The scale is normalized between +1 and −1 similar to AAP scale.

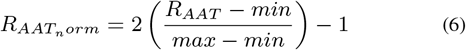

where max and min represent the maximum and minimum values among all the possible *R_AAT_* values before the normalization.

#### 2.2.6 K-mer representation

Segmentation of biological sequences into bag-of-or sequence-of-overlapping fixed length k-mers is one of the most favorable representations in bioinformatics research. K-mer representations are widely used in the areas of proteomics (Grabherr et al., 2011; Asgari and Mofrad, 2015), genomics (Jolma et al., 2013; Alipanahi et al., 2015), epigenomics (Awazu, 2016; Giancarlo et al., 2015), and metagenomics (Wood and Salzberg, 2014; Asgari et al., 2018).

In order to create a k-mer representation of a given protein sequence we divide it into overlapping subsequences of length *k* (k-mers). Subsequently, we represent the sequence as a frequency vector of all possible amino-acid k-mers (*vectorsize* = |20|^*k*^, where 20 is the number of amino-acids).

#### 2.2.7 Protvec sequence embeddings

Recently, in natural language processing (NLP), continuous vector representations of words known as word embeddings became very popular approach for word representation in the down-stream machine learning tasks. The general idea is to learn a vector representation of words in the course of neural probabilistic language modeling, and then use the learned representation as a general purpose representation of words in any NLP tasks. Language modeling is the task of assigning probability to a given sequence of words or predicting the next word given previous words. There are two main reasons for choosing language model-based representations:

i. language modeling is unsupervised and other information/metadata than the raw sentences are not needed. This lets us leverage the large amount of available text on the web for training a powerful representation,
ii. language modeling is a general purpose task, a representation that is relevant to language modeling is also relevant for syntactic and semantic similarities helping machine in the NLP tasks, e.g., machine translation, parsing, part-of-speech tagging or information retrieval.

Inspired by this idea, in a previous work we proposed distributed vector representations of biological sequence segments over k-mers, called biovector in general and ProtVec for proteins. We use the skip-gram neural network for this purpose. Skip-gram is a neural network with an objective analogous to a language modeling task (Mikolov et al., 2013; Bojanowski et al., 2016). However, instead of predicting the next word (or next k-mer) given the previous words, the task is to predict the surrounding words for a given word. We use large protein sequence databases (e.g., Swiss Prot) without any metadata for training a general purpose representation of protein k-mers. The objective of skip-gram neural network is to maximize the log-likelihood of observing contexts of k-mers in a window of *N* around it:

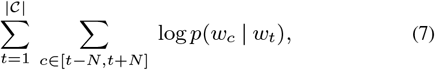

where *w_t_* is the current k-mer, *c*’s are the indices around index *t* in the window size of *N*. 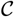 contains all existing k-mer contexts in the training data (e.g. all k-mer contexts exist in Swiss-Prot for all possible 3-mers). This likelihood is parameterized by k-mer representations (*v_t_*’s) and k-mer context representation (*v_c_*’s) in the skip-gram neural network:

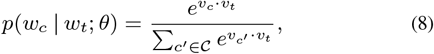

Since using all existing contexts in the above mentioned softmax function is computationally expensive, negative sampling is used in the training. After training the k-mer representations, in order to represent a given protein sequence we use summation embedding of existing k-mers in the sequence. Such representations have been proven helpful in protein function annotation tasks (Asgari and Mofrad, 2015; Zhou *et al.*, 2019).

### 2.3 Machine Learning Methods

After encoding the peptides as feature vectors, we use machine learning algorithms to classify the peptides as epitopes versus non-epitopes. For this binary classification, we employed two predictive models: Support Vector Machines (SVM) and Random Forest classifiers (RF). All the algorithms were implemented using the *Sklearn* package in Python.

Support vector machines (SVM) are a class of supervised machine learning methods usedforclassificationandregression. The SVM classifier learns a hyperplane that maximizes the geometric margin between positive and negative training data samples. When the training data are not linearly separable, a kernel function is used to map non-linearly separable data from the input space into a feature space. Here we used Radial Basis function (RBF) kernel. SVMs have been used extensively in linear epitope prediction (BCPred, LBTope, AAP etc.) (El-Manzalawy *et al.*, 2008; Singh *et al.*, 2013; Chen *et al.*, 2007) and they have also been employed for sequence-based prediction tasks (Zou *et al.*, 2013; Leslie *et al.*, 2002; Wu and Zhang, 2008). We used grid search to optimize the parameters *C* and γ over the range [1000 to 0.0001] with steps of power of 2.

Random forests are a class of supervised learning methods based on an ensemble of multiple decision trees which are mainly used for classification or regression. They make use of multiple decision trees and then take the mode of those to take a decision i.e. simply averaging multiple decision trees, trained on different parts of the same training set to reduce the variance. The random forest classifier we used in our method is an ensemble of 50 decision trees.

### 2.4 Performance Evaluation

We use five-fold cross validation where the dataset is randomly divided into five subsets with each containing an equal number of peptides. The subsets are then grouped into training and testing sets: four of the subsets are grouped into the training set and the remaining one is the testing set. The procedure is repeated five times with each subset used once for testing. The final prediction results are the average of the five testing results over the five-fold cross validation. We use several metrics to evaluate our prediction algorithms over the five-fold cross validation. Here, we calculate prediction accuracy *(ACC),* precision (Precision), recall/sensitivity (*S_n_*), F1 score (*F*1) and Mathews correlation coefficient *(MCC).* They are defined as follows:

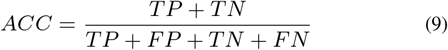

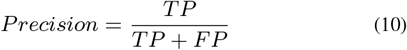

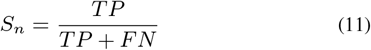

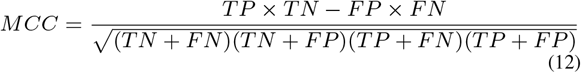

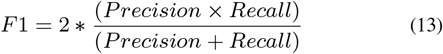

where TP, FP, TN, and FN are the numbers of true positives, false positives, true negatives, and false negatives, respectively. All these metrics are widely used to evaluate binary classification algorithms, but they all are threshold-dependent. If we vary the threshold we can vary the number of true positives at the cost of increasing false positives. We can plot the ROC (receiver operating characteristic) curve by plotting the true positive rate as a function of the false positive rate and it shows the performance of the classifier over all possible thresholds.. The area under the ROC curve (AUC) can be used to compare the curves and it can be interpreted as the probability that a randomly chosen positive example will be ranked higher than a randomly chosen negative example.

## 3 Results

We trained and evaluated our method with five-fold cross validation on the BCPred dataset. We here describe the results of using different features and report the performance of our method when tested on multiple datasets.

### 3.1 Discriminative power of individual features

We used several feature sets for our machine-learning models and investigated which feature set has the best discriminative power when used independently. We tested the SVM model on the BCPred dataset with five-fold cross validation while taking individual feature sets (Table 2). All of the machine-learning models were trained on the BCPred dataset with five-fold cross validation, and C and *γ* parameters for the RBF kernel were optimized with grid-search, and performance was averaged over the hold-out folds. CTD feature had an acuracy of 61% while the amino acid composition scales performed quite similarly with their accuracy ranging from 63% (AAC) to 65% (DPC). With k-mer representation, higher value of *k* resulted in a better accuracy (highest being 69.9% with *k*=4) and ProtVec had an accuracy of 70%. The AAP antigenicity scale had an accuracy of 68.55% and AAT antigenicity scale had the highest accuracy of 78.67%.

**Table 1.** Summary of datasets used in the benchmarking of our approach. The datasets are available at https://github.com/hzi-bifo/epitope-prediction-paper

| Dataset | Original method | Epitopes | Non-epitopes | Source |
| --- | --- | --- | --- | --- |
| BCPred | BCPred | 700 | 700 | BciPep |
| ABCPred16 | ABCPred | 701 | 701 | BciPep |
| Chen | AAP | 872 | 872 | BciPep |
| Blind387 | ABCPred | 187 | 200 | BciPep |
| LBTope_fixed_nr | LBTope | 7824 | 7853 | IEDB |
| Viral | New | 4432 | 8460 | IEDB |

**Table 2.** Comparison of individual feature sets in the epitope prediction in terms of ROC_AUC, accuracy, and F1 score validated with a 5-fold cross-validation over the BCPred dataset.

| Feature | ROC_AUC | Accuracy | F1 |
| --- | --- | --- | --- |
| AAC | 0.683 | 63.13 | 0.611 |
| DPC | 0.701 | 65.83 | 0.593 |
| CTD | 0.645 | 61.20 | 0.584 |
| AAP(average) | 0.708 | 69.04 | 0.692 |
| AAP | 0.733 | 68.55 | 0.680 |
| AAT | 0.878 | 78.67 | 0.782 |
| k-mer (k=1) | 0.678 | 63.12 | 0.599 |
| k-mer (k=2) | 0.700 | 66.26 | 0.613 |
| k-mer (k=3) | 0.721 | 68.71 | 0.639 |
| k-mer (k=4) | 0.746 | 69.93 | 0.660 |
| Protvec | 0.740 | 70.02 | 0.693 |

### 3.2 Discriminative power of the combination of features

We next investigated combinations of different feature types to optimize the performance of our ML model (Fig 1). We looked at AAC, DPC, AAP, AAT, 4-mers and ProtVec feature sets (Table 3). We considered some combinations of these features and then combined features based on the feature type. The combination of AAC and DPC achieved an accuracy of 66.20%, which was slightly higher thantheindividualaccuracyof the AAC or DPC features. On combining the composition based features (AAP, AAT and AAC) and sequence representation based features (Protvec), the highest accuracy of 81.31% was achieved and this was selected as our new method (EpitopeVec).

**Fig. 1:**
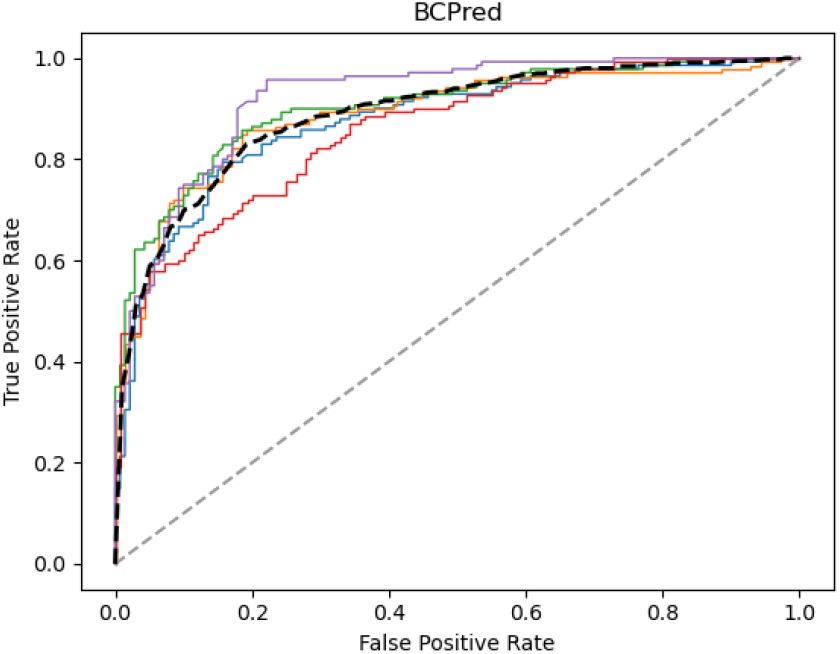
ROC-curve of epitope prediction in a five fold cross-validation on the BCPred dataset, where the mean is shown bold/dashed line and the random performance (ROC_AUC=0.5) is shown in gray/dashed.

**Table 3.**
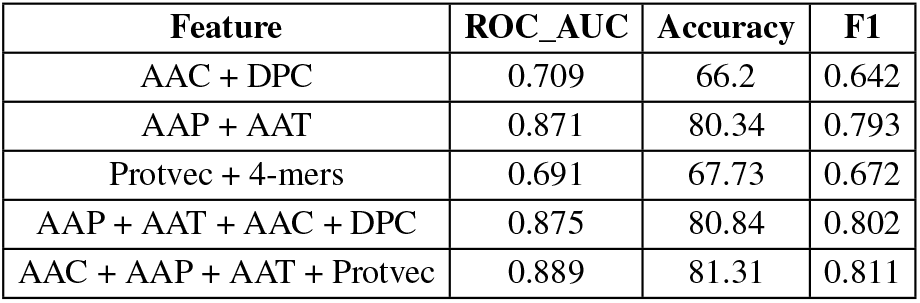
Comparison of feature combinations in the epitope prediction in terms of ROC_AUC, accuracy, and F1 score validated with a 5-fold cross-validation over the BCPred dataset.

### 3.3 Evaluation on the BCPred dataset

We compared our method to the results of six other methods on the BCPred dataset. On five-fold cross-validation, our method performed the best in all the metrics for this dataset with an average accuracy of 81.31% (Table 4) and ROC of 0.889 (Fig 2). It was substantially better than the original BCPred method (El-Manzalawy et al., 2008), which achieved an accuracy of 67.9% (13.41 % higher and 19.7% improvement on the original). The performance of methods trained on the IEDB datasets, SVMTrip (AUC = 0.572) and LBtope (accuracy = 51.57%), was reported to be poor (Singh et al., 2013) (Yao et al., 2012). The performance of methods trained on datasets from BCIPep database (BCPred, AAP, ABCPred) was reported to be between 64% to 71%.

**Fig. 2:**
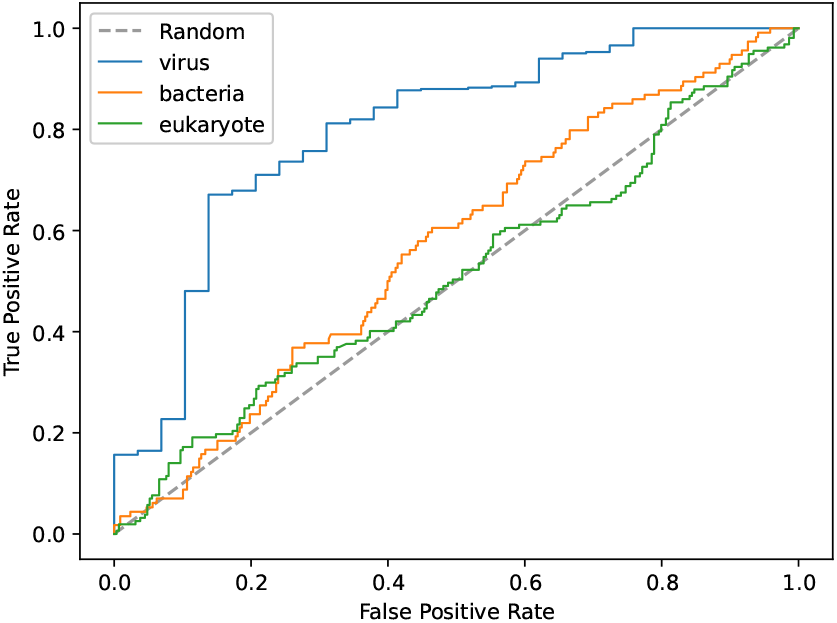
Area under the ROC curve for each antigen type in the BCPred set, where the EpitopeVec-viral predictor is utilized as the predictive model.

### 3.4 Evaluation on the Chen dataset

The Chen method (AAP method) made use of a novel scale, called the amino acid pair (AAP) antigenicity scale, which proposed that certain AAPs are favored in epitope regions. It is the first method to incorporate the differences between epitopes and non-epitopes by making use of AAP antigenicity scale as a discriminative feature. We achieved an accuracy of 88.30% (Table 4) on testing our method on the Chen dataset, which is 10% higher than the next best method Cobe-Pro(78%) and 16% higher than the original AAP method (Chen et al., 2007). This dataset is not homology reduced, which explains why we found a higher accuracy than for the BCPred dataset. As expected, the performance of the newer methods like LBTope and SVMTrip on this dataset was reported to be poor (accuracy around 53.33%) because they are trained on IEDB-derived datasets (Yao et al., 2012). The Chen dataset is also a BCIPep derived dataset and therefore the performance of BCIPep trained methods on this dataset was similar to that on BCPred set.

### 3.5 Evaluation on the ABCPred dataset

The original ABCPred method is based on a recurrent neural network with an input vector of 16 residues using a sparse binary encoding and it is one of the first methods to use machine learning for linear epitope prediction. We tested our method (trained on BCPred) on ABCPred16 (length of 16 amino acids) and achieved a higher accuracy of 86.50% (20.57% higher and 31.2 % improvement on the original) in comparison with the original ABCPred method (65.93%) (table 4). The performance of methods such as AAP, BCPred, ABCPred that were trained on BciPep-derived datasets was reported to be between 65% to 73% (Chen et al., 2007; El-Manzalawy et al., 2008). The performance of LBTope (trained on IEDB-derived dataset) was reported to be 57.90% (Singh et al., 2013). We also tested the method on the Blind387 set that was published along with ABCPred (Saha and Raghava, 2006). In this dataset the length of the epitopes is not fixed and it varies from 5 to 30 amino acids. Our method was trained on the BCPred set and then tested on this Blind387 set. Our method obtained an accuracy of 72.42%, outperforming accuracies previously reported by AAP (64.60%), ABCPred (66.41%) and BCPred (65.89%) (Chen et al., 2007; Saha and Raghava, 2006; El-Manzalawy et al., 2008).

### 3.6 Evaluation on the LBTope dataset

LBtope is one of the first methods that made use of datasets compiled from IEDB dataset. Our method achieved an accuracy of 52.98% in comparison with the best accuracy of BCPred (52.56%) in cross-dataset testing. The substantial reduction in the performance was observed because our method was trained on the BCPred dataset that was compiled from the BciPep database and LBTope was compiled from the IEDB. To compare our method directly with the LBTope method, we also trained our method on the LBTope set and reported the performance using five-fold cross validation (Table 4). The accuracy with five-fold cross-validation on the LBTope set using EpitopeVec was 11% higher (17% improvement over original LBTope method) than the best accuracy reported in the original LBTope study (Singh et al., 2013).

### 3.7 Improving epitope-prediction accuracy using domain-specific datasets

The substantial performance difference we observed between cross-testing (accuracy of 53%) and 5-fold cross-validation on the BCPRED dataset (acuracy of 81.31%) (Table 4) from the BciPep database and the LBTOPE dataset (accuracy of 76.01%) from the IEDB database indicates a lack of generalization ability of the classifiers, potentially due to the fundamental differences in the nature of the underlying data (Odorico and Pellequer, 2003). Previously published methods like BCPred and LBTope also reported poor accuracy (51% – 52%) when they were tested on independent datasets derived from databases different than their training sets. When we looked at the Pearson correlation of the AAT scale (the most discriminative feature in our method) derived from BciPep-sets and IEDB-sets, it was relatively low (0.41), indicating large differences in the composition of the epitopic peptides from both databases. When analyzing the composition of these databases, we found that BciPep has a strong bias towards viruses (80% peptides are from viral antigens). To investigate this further, we created a dataset of viral epitopes (with experimentally verified positive and negative epitopes) from the IEDB dataset. We trained a viral predictor on this dataset with EpitopeVec, optimized the C and *γ* parameters for the RBF kernel and cross-tested it on BCPred set. BCPred is composed of antigens of all types, i.e. viral, bacterial, eukaryotic, so we looked at antigen-type specific performance measures. The accuracy for viral peptides (72.57%) was substantially higher than that for bacterial (61.73%) and eukaryotic (56.73%) peptides (Table 4). The area under the ROC curves is also substantially higher for viral peptides (0.799) than for bacterial (0.569) and eukaryotic peptides (0.512) (Fig 3). This indicates that properties distinguishing between epitopic and non-epitopic peptides are specific to antigen type and creating a general purpose classifier that performs well on any antigen type is difficult. The performance of the viral predictor substantially improves on previous methods and it could be used to predict linear viral BCE’s.

**Table 4.** Comprehensive benchmarking of different linear BCE predictors on different data sets, where EpitopeVec (our approach) is compared with the existing methods. The empty gray cells show the cases where the corresponding metrics in the software were not available and we only could include the score reported by the original paper.

| Method | ROC_AUC | Accuracy | Precision + | Precision - | Recall + | Recall - | F1 | MCC |
| --- | --- | --- | --- | --- | --- | --- | --- | --- |
| **BCPred dataset** |  |  |  |  |  |  |  |  |
| *ABCPred*<sup>1</sup> | 0.643 |  |  |  |  |  |  |  |
| *AAP*<sup>2</sup> | 0.7 | 64.05 |  |  |  |  |  |  |
| *BCPred*<sup>3</sup> | 0.758 | 67.9 |  |  |  |  |  | 0.36 |
| *Cobe - Pro*<sup>4</sup> | 0.768 | 71.4 |  |  |  |  |  |  |
| *SVMTrip*<sup>5</sup> | 0.667 |  |  |  |  |  | 0.572 |  |
| *LBTope*<sup>6</sup> |  | 51.57 |  |  |  |  |  |  |
| *EpitopeVec* | **0.889** | **81.31** | **0.816** | **0.811** | **0.807** | **0.819** | **0.811** | **0.627** |
| **Chen dataset** |  |  |  |  |  |  |  |  |
| *AAP*<sup>2</sup> | 0.7 | 71.09 |  |  |  |  |  | 0.366 |
| *AAP + scales*<sup>2</sup> |  | 72.54 |  |  |  |  |  | 0.404 |
| *APCPred*<sup>7</sup> | 0.809 | 72.94 |  |  |  |  |  | 0.36 |
| *Cobe - Pro*<sup>4</sup> | 0.829 | 78 |  |  |  |  |  |  |
| *SVMTrip*<sup>5</sup> | 0.667 |  |  |  |  |  | 0.59 |  |
| *LBTope*<sup>6</sup> |  | 53.33 |  |  |  |  |  |  |
| *EpitopeVec* | **0.958** | **88.30** | **0.849** | **0.932** | **0.925** | **0.833** | **0.889** | **0.770** |
| **ABCPred dataset** |  |  |  |  |  |  |  |  |
| *AAP*<sup>2</sup> | 0.782 | 73.14 |  |  |  |  |  | 0.518 |
| *ABCPred*<sup>1</sup> |  | 65.93 |  |  |  |  |  | 0.3187 |
| *BCPred*<sup>3</sup> | 0.758 | 67.90 |  |  |  |  |  | 0.36 |
| *APCPred*<sup>7</sup> | 0.794 | 73.00 |  |  |  |  |  |  |
| *LBTope*<sup>6</sup> |  | 57.90 |  |  |  |  |  |  |
| *EpitopeVec* | **0.928** | **86.50** | **0.852** | **0.882** | **0.879** | **0.847** | **0.867** | **0.730** |
| **Blind387 dataset** |  |  |  |  |  |  |  |  |
| *AAP*<sup>2</sup> | 0.689 | 64.60 |  |  |  |  |  | 0.292 |
| *ABCPred*<sup>1</sup> |  | 66.41 |  |  |  |  |  | 0.3187 |
| *BCPred*<sup>3</sup> | 0.699 | 65.89 |  |  |  |  |  | 0.318 |
| *EpitopeVec* | **0.781** | **72.42** | **0.760** | **0.626** | **0.701** | **0.816** | **0.686** | **0.451** |
| **LBTope dataset** |  |  |  |  |  |  |  |  |
| *LBTope*<sup>6</sup> | 0.69 | 64.86 |  |  |  |  |  |  |
| *BCPred*<sup>3</sup> |  | 52.56 |  |  |  |  |  |  |
| *EpitopeVec(BCPred)\** | 0.548 | 52.98 | 0.550 | 0.321 | 0.522 | 0.738 | 0.508 | 0.065 |
| *EpitopeVec(LBTope)<sup>§</sup>* | **0.841** | **76.01** | **0.757** | **0.763** | **0.765** | **0.755** | **0.761** | **0.521** |
| **Viral dataset** |  |  |  |  |  |  |  |  |
| *EpitopeVec(Viral)* | 0.875 | 79.73 | 0.718 | 0.843 | 0.698 | 0.67 | 0.850 | 0.554 |
| *EpitopeVec(onBCPred - viral)* | 0.799 | 72.57 | 0.975 | 0.172 | 0.723 | 0.759 | 0.830 | 0.266 |
| *EpitopeVec(onBCPred - bacteria)* | 0.569 | 61.73 | 0.297 | 0.769 | 0.377 | 0.700 | 0.332 | 0.070 |
| *EpitopeVec(onBCPred - eukaryote)* | 0.512 | 56.73 | 0.378 | 0.660 | 0.350 | 0.685 | 0.363 | 0.036 |

1 : Saha and Raghava (2006), 2 : Chen *et al.* (2007), 3 : El-Manzalawy *et al.* (2008), 4 : Sweredoski and Baldi (2009), 5 : Yao *et al.* (2012), 6 : Singh *et al.* (2013), 7 : Shen *et al.* (2015)
\* : Trained on BCPred set and tested on LBTope set, §: Five-fold cross validation on LBTope set

### 3.8 Predicting epitopes for SARS-CoV-1 and SARS-CoV-2 viruses

We tested our viral predictor on peptides suggested in (Grifoni et al., 2020) for SARS viruses. They compiled a set of 10 linear peptides for SARS-CoV-1 proteins from the experimentally verified epitopes in IEDB and then mapped these 10 regions to the SARS-CoV-2 proteins. As SARS-CoV has a high sequence similarity to SARS-CoV, they suggested that the corresponding regions on SARS-CoV could be potential linear b-cell epitopes. We tested the suggested peptides using our method to assess how many were classified as epitopes. We predicted 7 out of 10 peptides as epitopes for SARS-CoV and 7 out of 10 for SARS-CoV-2 (Table 5). This demonstrates the biological viability of our

**Table 5.**
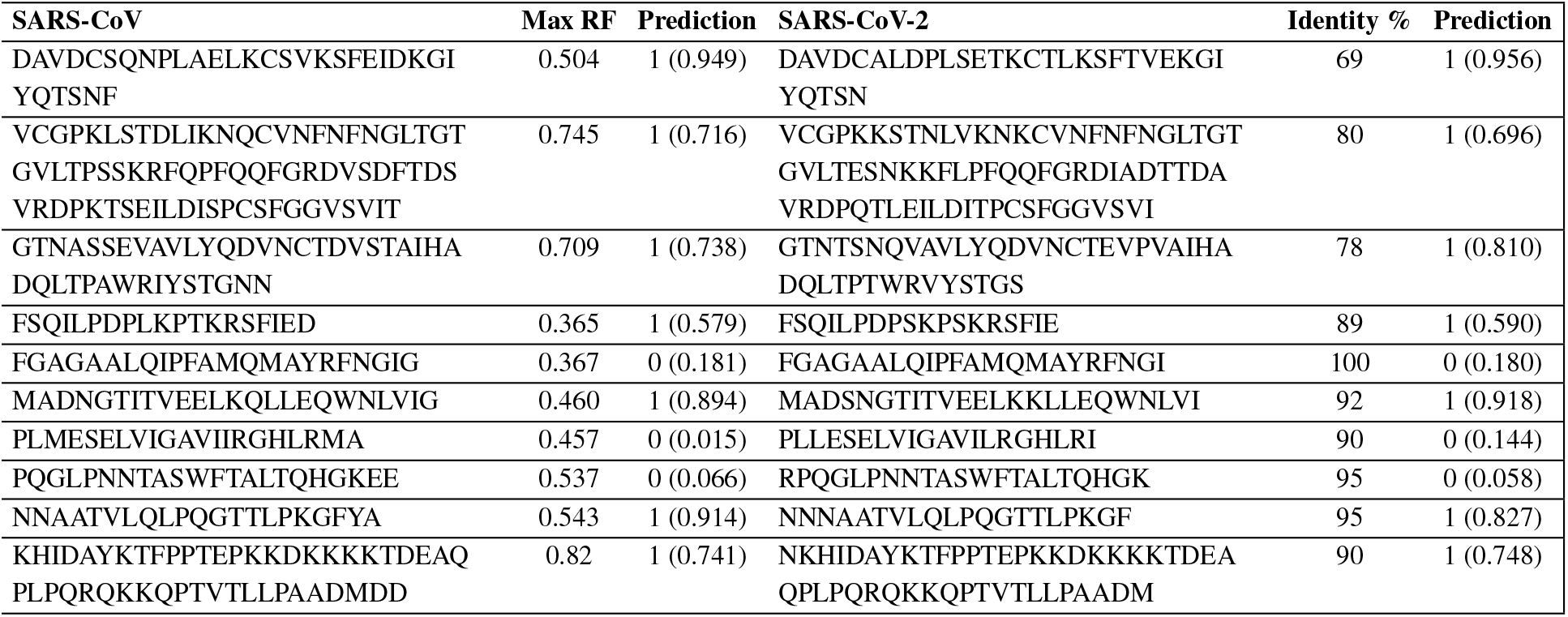
Predictions on Corona virus peptides by EpitopeVec-viral

## 4 Discussion and Conclusion

Here, we describe a computational framework for predicting linear B-cell epitopes using a combination of amino acid features, modified antigenicity scales (AAT), and the ProtVec embedding of peptides as the feature vector.

The use of the AAT scale has already been explored by the SVMTrip method. However, in our implementation, we proposed the scale’s negative logarithm before normalization, improving its discriminative power. Sequence-based embeddings have been used successfully in protein functional/structural annotations tasks previously such as secondary structure prediction (Li and Yu, 2016; Asgari et al., 2019a), point mutations (Zhou et al., 2020), protein function prediction (Asgari and Mofrad, 2015; Zhou et al., 2019; Bonetta and Valentino, 2020), and predicting structural motifs (Liu et al., 2018). In this paper, we proposed the use of ProtVec embeddings and k-mers for linear BCE prediction improving state-of-the-art performance on different datasets.

To assess the generalizability of the EpitopeVec model, we performed a benchmarking of different predictive models over existing datasets for BCE prediction. Our benchmarking of different predictive models over the BCPred dataset showed that EpitopeVec outperformed the state-of-the-art approach by 10% accuracy in 5-fold cross-validation, which is also 13.41% higher than that of the original BCPred method. We evaluated the trained model over the BCPred dataset directly over BciPep, Blind387, and IEDB (LBTope and SVMTrip) datasets. EpitopeVec could successfully generalize over the BciPep and Blind387 datasets, achieving state-of-the-art accuracy (72% to 88%). However, the EpitopeVec accuracy (pretrained over the BCPred dataset) lowered to 53% on the IEDB dataset. Further investigations confirmed that although the pre-trained model over the BCPred dataset could not generalize over IEDB, the 5-fold cross-validation on the same dataset obtains an accuracy of 76%, which is 11% higher than that of the original LBTOPE method. (Singh et al., 2013).

Our benchmarking results indicate that the difference between the cross-validation performance on the IEDB and the cross-testing of a pre-trained model (trained on BCPred) tested on the IEDB roots in the different composition of residues in the +ve/-ve sets of these datasets. This observation suggests that the residues properties distinguishing epitopes from non-epitopes are not generalizable across antigens types and rely on certain factors (e.g., whether we are dealing with a viral, bacterial, or a fungal antigen). The apparent difficulties of creating an accurate general-purpose linear BCE predictor proposes a future direction of creating specialized predictors for specific antigenic types. We trained a linear viral B-cell epitope predictor on a viral dataset separately in favor of this conclusion. On cross-testing on the BCPred dataset, the viral predictor performed substantially better on viral peptides compared to that on bacterial and prokaryotic peptides. We were able to predict 7 out of 10 epitopes correctly for SARS-CoV-1, which demonstrates good biological viability of our viral predictor. This predictive model can be used on viral proteins and potentially help experimentalists design new peptide-based vaccines.

In general, the prediction of linear BCEs is considered to be more challenging than T-cell epitopes (Soria-Guerra *et al.*, 2015), as the length of BCEs is not fixed (can vary from 5 to 30 residues). To overcome this problem, we made the predictions over the fixed-length epitopes of length 20 (except a few variable-length datasets) covering most functional epitopes (Kringelum *et al.*, 2013). As many naturally-occurring BCEs are conformational (Sivalingam and Shepherd, 2012) and not contiguous stretches in the primary protein sequence, the inclusion of tertiary structure information and prediction of conformational epitopes could be the next step in the development of BCE predictive models.

Another challenge for the ML-based prediction of epitopes is creating reliable negative samples. Most datasets introduce a random sub-sample of peptides from SWISSPROT as their negative samples, and some have taken the region from the antigen sequence not reported as epitope to be non-epitope. However, as the available experimental data are not complete, some of these non-epitope residues might be epitope residues binding to different antibodies. In addition, there could be multiple epitopes in an antigen sequence not discovered when compiling datasets from neutralization experiments, as most of the experiments only test against a particular antibody. Incorrect negative samples increase the false-negative rate of the predictive model and subsequently lower the prediction performances. For future development, large, non-redundant, and experimentally well-characterized datasets could be compiled and standardized for the training and the evaluation of BCE predictive models.

## Acknowledgements

The authors would like to thank Tzu-Hao Kuo for his valuable comments and helping out with testing the code.

## References

Alipanahi, B. et al. (2015). Predicting the sequence specificities of dna-and rna-binding proteins by deep learning. Nat. Biotechnol., 33(8), 831–838.

Alix, A. J. (1999). Predictive estimation of protein linear epitopes by using the program PEOPLE. In Vaccine, volume 18, pages 311–314. Elsevier.

Asgari, E. (2019). Life Language Processing: Deep Learning-based Language-agnostic Processing of Proteomics, Genomics/Metagenomics, and Human Languages. Ph.D. thesis, UC Berkeley.

Asgari, E. and Mofrad, M. R. (2015). Continuous distributed representation of biological sequences for deep proteomics and genomics. PloS One, 10(11), e0141287.

Asgari, E. et al. (2018). Micropheno: predicting environments and host phenotypes from 16s rrna gene sequencing using a k-mer based representation of shallow sub-samples. Bioinformatics, 34(13), i32–i42.

Asgari, E. et al. (2019a). Deepprime2sec: Deep learning for protein secondary structure prediction from the primary sequences. bioRxiv, page 705426.

Asgari, E. et al. (2019b). Probabilistic variable-length segmentation of protein sequences for discriminative motif discovery (dimotif) and sequence embedding (protvecx). Scientific reports, 9(1), 1–16.

Awazu, A. (2016). Prediction of nucleosome positioning by the incorporation of frequencies and distributions of three different nucleotide segment lengths into a general pseudo k-tuple nucleotide composition. Bioinformatics, 33(1), 42–48.

Bairoch, A. (2000). The SWISS-PROT protein sequence database and its supplement TrEMBL in 2000. Nucleic Acids Research.

Barlow, D. J. et al. (1986). Continuous and discontinuous protein antigenic determinants. Nature, 322(6081), 747–748.

Blythe, M. J. and Flower, D. R. (2009). Benchmarking B cell epitope prediction: Underperformance of existing methods. Protein Science, 14(1), 246–248.

Bojanowski, P. et al. (2016). Enriching word vectors with subword information. arXiv preprint arXiv:1607.04606.

Bonetta, R. and Valentino, G. (2020). Machine learning techniques for protein function prediction.

Bryson, C. J. et al. (2010). Prediction of immunogenicity of therapeutic proteins: Validity of computational tools.

Cao, D. S. et al. (2013). PyDPI: Freely available python package for chemoinformatics, bioinformatics, and chemogenomics studies. Journal of Chemical Information and Modeling.

Chen, J. et al. (2007). Prediction of linear B-cell epitopes using amino acid pair antigenicity scale. Amino Acids.

Dubchak, I. et al. (1995). Prediction of protein folding class using global description of amino acid sequence. Proceedings of the National Academy of Sciences of the United States of America.

Dudek, N. L. et al. (2010). Epitope discovery and their use in peptide based vaccines. Curr. Pharm. Des., 16(28), 3149–3157.

El-Manzalawy, Y. et al. (2008). Predicting linear B-cell epitopes using string kernels. Journal of Molecular Recognition.

Emini, E. A. et al. (1985). Induction of hepatitis A virus-neutralizing antibody by a virus-specific synthetic peptide. Journal of Virology.

Flower, D. R. (2007). Immunoinformatics. Predicting immunogenicity in silico. Preface.

Giancarlo, R. et al. (2015). Epigenomic k-mer dictionaries: shedding light on how sequence composition influences in vivo nucleosome positioning. Bioinformatics, 31(18), 2939–2946.

Grabherr, M. G. et al. (2011). Full-length transcriptome assembly from rna-seq data without a reference genome. Nat. Biotechnol., 29(7), 644–652.

Grifoni, A. et al. (2020). A Sequence Homology and Bioinformatic Approach Can Predict Candidate Targets for Immune Responses to SARS-CoV-2. Cell Host and Microbe.

Hancock, D. C. and O’Reilly, N. J. (2005). Synthetic peptides as antigens for antibody production. Methods in molecular biology (Clifton, N.J.).

Huang, Y. et al. (2010). CD-HIT Suite: A web server for clustering and comparing biological sequences. Bioinformatics.

Janeway, C. (2012). immunobiology, 5th ed.

Jolma, A. et al. (2013). Dna-binding specificities of human transcription factors. Cell, 152(1-2), 327–339.

Karplus, P. A. and Schulz, G. E. (1985). Prediction of chain flexibility in proteins - A tool for the selection of peptide antigens. Naturwissenschaften.

Kolaskar, A. S. and Tongaonkar, P. C. (1990). A semi-empirical method for prediction of antigenic determinants on protein antigens. FEBS Letters.

Kringelum, J. V. et al. (2013). Structural analysis of B-cell epitopes in antibody: Protein complexes. Molecular Immunology.

Larsen, J. et al. (2006). Improved method for predicting linear B-cell epitopes. Immunome Research, 2(1), 2.

Leslie, C. et al. (2002). The spectrum kernel: a string kernel for SVM protein classification. Pacific Symposium on Biocomputing. Pacific Symposium on Biocomputing.

Levitt, M. (1976). A simplified representation of protein conformations for rapid simulation of protein folding. Journal of Molecular Biology.

Li, Z. and Yu, Y. (2016). Protein secondary structure prediction using cascaded convolutional and recurrent neural networks. In IJCAI International Joint Conference on Artificial Intelligence.

Liu, Y. et al. (2018). Learning structural motif representations for efficient protein structure search. In Bioinformatics.

Mikolov, T. et al. (2013). Distributed representations of words and phrases and their compositionality. In Advances in Neural Information Processing Systems, pages 3111–3119.

Noya, O. et al. (2005). Immunodiagnosis of Parasitic Diseases with Synthetic Peptides. Current Protein & Peptide Science.

Odorico, M. and Pellequer, J. L. (2003). BEPITOPE: Predicting the location of continuous epitopes and patterns in proteins. Journal of Molecular Recognition, 16(1), 20–22.

Pellequer, J. L. and Westhof, E. (1993). PREDITOP: A program for antigenicity prediction. Journal of Molecular Graphics, 11(3), 204–210.

Pellequer, J. L. et al. (1991). Predicting location of continuous epitopes in proteins from their primary structures. Methods in Enzymology.

Saha, S. and Raghava, G. P. (2006). Prediction of continuous B-cell epitopes in an antigen using recurrent neural network. Proteins: Structure, Function and Genetics.

Saha, S. et al. (2005). Bcipep: A database of B-cell epitopes. BMC Genomics.

Shen, W. et al. (2015). Predicting linear B-cell epitopes using amino acid anchoring pair composition. BioData Mining.

Singh, H. et al. (2013). Improved Method for Linear B-Cell Epitope Prediction Using Antigen’s Primary Sequence. PLoS ONE.

Sivalingam, G. N. and Shepherd, A. J. (2012). An analysis of B-cell epitope discontinuity. Molecular Immunology.

Soria-Guerra, R. E. et al. (2015). An overview of bioinformatics tools for epitope prediction: Implications on vaccine development.

Sweredoski, M. J. and Baldi, P. (2009). COBEpro: A novel system for predicting continuous B-cell epitopes. Protein Engineering, Design and Selection.

Vita, R. et al. (2009). The Immune Epitope Database 2.0. Nucleic Acids Research.

Wood, D. E. and Salzberg, S. L. (2014). Kraken: Ultrafast metagenomic sequence classification using exact alignments. Genome Biol., 15(3), R46.

Wu, S. and Zhang, Y. (2008). A comprehensive assessment of sequencebased and template-based methods for protein contact prediction. Bioinformatics.

Yang, X. and Yu, X. (2009). An introduction to epitope prediction methods and software.

Yao, B. et al. (2012). SVMTriP: A Method to Predict Antigenic Epitopes Using Support Vector Machine to Integrate Tri-Peptide Similarity and Propensity. PLoS ONE, 7(9), e45152.

Zhou, G. et al. (2020). Mutation effect estimation on protein-protein interactions using deep contextualized representation learning. NAR Genomics and Bioinformatics.

Zhou, N. et al. (2019). The cafa challenge reports improved protein function prediction and new functional annotations for hundreds of genes through experimental screens. Genome biology, 20(1), 1–23.

Zou, C. et al. (2013). An improved sequence based prediction protocol for DNA-binding proteins using SVM and comprehensive feature analysis. BMC Bioinformatics.

